# Parting the cellular sea: electrotaxis evoked directional separation of co-cultured keratinocytes and fibroblasts

**DOI:** 10.1101/2023.04.13.536575

**Authors:** José Leal, Sebastian Shaner, Nicole Jedrusik, Anna Savelyeva, Maria Asplund

## Abstract

Bioelectric communication plays a significant role in several cellular processes and biological mechanisms, such as division, differentiation, migration, cancer metastasis, or wound healing. The flow of ions through cellular walls and the gradients generated thereby evoke this signaling as electric fields (EFs) form across membranes, and their changes act as cues for cells. An EF is natively generated towards the wound center during epithelial wound healing, aiming to align and guide cell migration, particularly of keratinocytes, fibroblasts, and macrophages. While this phenomenon, known as electrotaxis, has been extensively investigated across many cell types, it is typically explored one cell type at a time, which does not accurately represent cellular interactions during complex biological processes. Here we show the co-cultured electrotaxis of epithelial keratinocytes and fibroblasts with a salt-bridgeless microfluidic approach for the first time. The electrotactic response of these cells was first assessed in mono-culture to establish a baseline, resulting in a characteristic anodic migration for keratinocytes and cathodic for fibroblasts. Both cell types retained their electrotactic properties in co-culture leading to clear cellular partition. The methods leveraged herein can pave the way for future co-culture electrotaxis experiments where the concurrent influence of cell lines can be thoroughly investigated.

## 1. Introduction

Bioelectricity plays a role in all mammalian somatic cells, not just excitable ones. The main bioelectric difference between excitable and non-excitable cells is the time scale: milliseconds for the former and minutes to days for the latter.^1^ While action potentials are how information is relayed in excitable cells, non-excitable cells depend on gap junctions to pass along bioelectric information. A canonical example is found within the epidermis during wound healing. A gradient of ions from the outermost layer (apical) to the innermost layer (basal) of the epidermis generates a transepithelial potential (TEP). When the skin and TEP are broken, a new ionic gradient is formed, with the current pointing toward the center of the newly-formed wound. This endogenous electric field (EF) is crucial in wound healing. It acts as a bioelectric guidance cue for nearly all cells responsible for cleanup, repair, and remodeling.

The directed migration of cells along or against an EF’s direction is called electrotaxis or galvanotaxis. This phenomenon occurs throughout the body, from wound healing to cancer invasion,^2,3^ and has been extensively investigated *in vitro* in over 30 cell types (both non-excitable and excitable).^4,5^ Several biochemical mechanisms have been reported to be directly involved and responsible for electrotaxis: signaling pathways (e.g., phosphatidylinositol-3-OH kinase (P3IK), mitogen-activated protein kinase (MAPK)),^6,7^ voltage-gated ion channels (e.g., sodium- and calcium-selective),^8,9^ and growth factors (e.g., epidermal, vascular endothelial).^10^ While these relate to specific cell types, more global bioelectric mechanisms have also been identified. These impart asymmetrical mechanical intracellular forces, and sequential intracellular pathway activation cascades through spatial and polar redistribution of charged cell membrane receptors (e.g., ion channels, integrins) through electrophoretic or electroosmotic forces.^11–14^

The interplay between bioelectric and biochemical driving factors during electrotaxis likely overlaps across cell types and species to varying degrees. The culmination of these factors gives rise to a direct electrotactic directionality response across all the investigated cell types. The vast majority of cells migrate toward the more negatively-charged electrode (i.e., cathode); however, some cell types migrate toward the anode.^15^ This electrotactic phenotype duality is also a characteristic of the cells partaking in the wound-healing process. Dermal fibroblasts migrate anodically toward the wound’s edges to restore the extracellular matrix (ECM) foundation from the inside out, while keratinocytes migrate to the wound’s center cathodically to rebuild the epidermis on the newly built ECM from the outside in. This differing electrotactic response for keratinocytes and fibroblasts has been shown *in vitro* to be strongly regulated by PI3K signaling. Furthermore, these cells show an electrotactic response at different EF strengths. Keratinocytes exhibit directed migration at 100 mV mm^-1^ after 1 h, while fibroblasts require either longer times (3 h at 100 mV mm^-1^) or higher EFs (400 mV mm^-1^).^6^ While this bipolar electrotactic response has been observed *in vivo* for keratinocytes and fibroblasts, it has not yet been concurrently observed in an *in vitro* live-cell imaging setting.^16–18^

To investigate how electrotaxis fundamentally works, one must recreate the endogenous driving force in biology, meaning a constant EF over a defined time frame, and observe how these fields affect cellular mechanisms, morphology, metabolism, and movement. Microfluidic devices can provide precise setups for electrotaxis research by controlling the distribution and tuning of electric fields (EF) in cell-containing microchannels. ^19,20^ However, there are some challenges associated with cell seeding, such as restricted volume and low surface area. ^21^ A technique that allows spot-cast seeding into laser-cut microchannels with subsequent device assembly previously developed by our group is leveraged here to overcome these difficulties. ^22^ The electrodes employed for DC stimulation, such as silver/silver chloride (Ag/AgCl) or platinum (Pt), can produce cytotoxic chemical products and require careful experimental design prior to experimentation (i.e., use of salt bridges). ^23–28^ To address this issue, we previously developed salt-bridgeless systems, which rely on metal or carbon electrodes coated with a supercapacitive conducting polymer or hydrogel based on PEDOT:PSS, which we leverage in this work. ^29,30^ These electrodes have outstanding biocompatibility and stable DC stimulation capabilities with cultured cells, providing a streamlined alternative for faster *in vitro* electrotaxis experimentation with potential for future *in vivo* applications.^31^

In this study, we combine these technological developments to address two questions: **(1)** Will mono-cultured keratinocytes and fibroblasts stimulated in salt-bridgeless microfluidic devices electrotactically concur with findings using conventional salt-bridge systems? **(2)** Will co-cultured keratinocytes and fibroblasts exhibit the same electrotactic behavior as their mono-cultured counterparts? (**Figure 1**)

**Figure 1:**
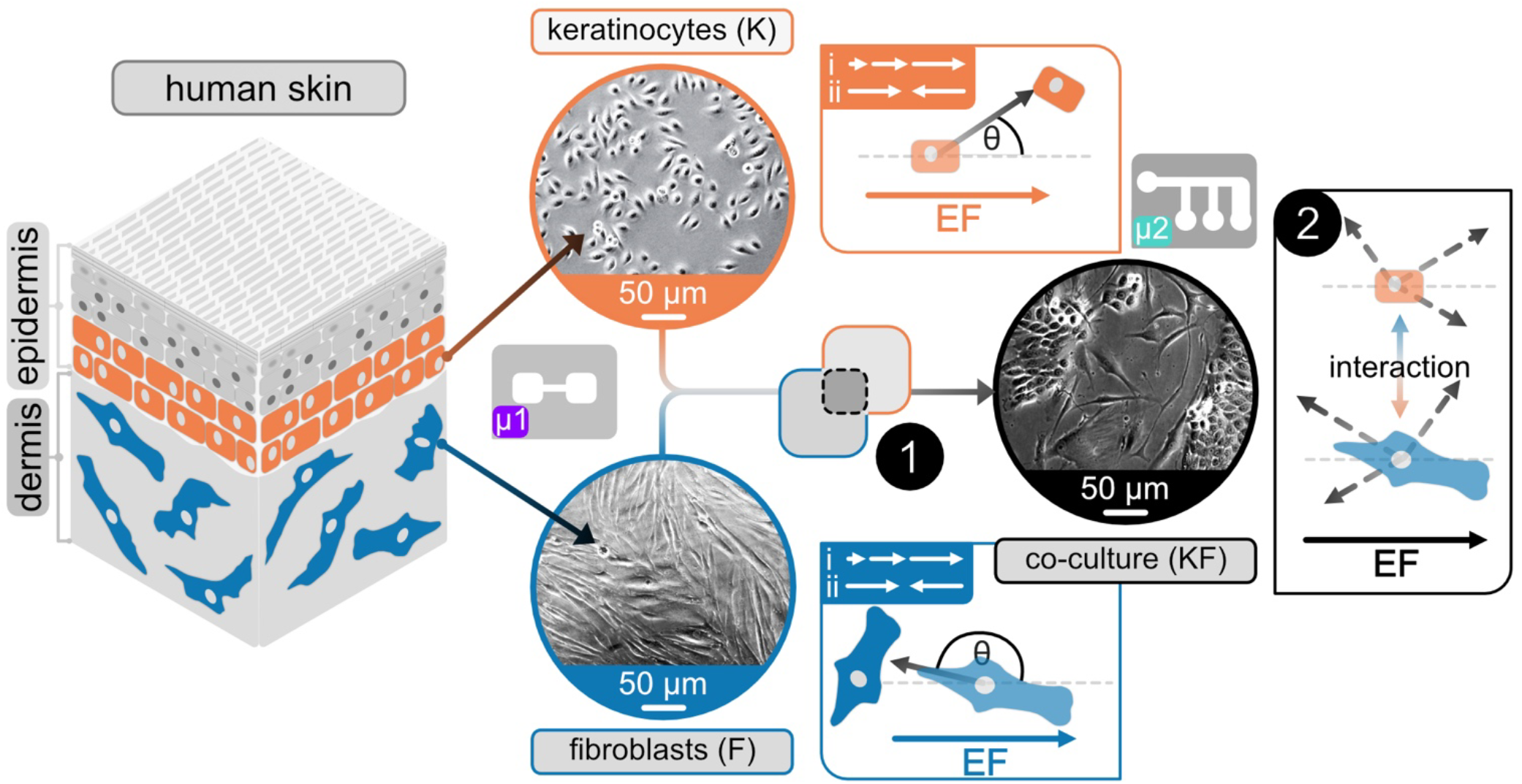
Visual representation of hypothesis. Keratinocytes and fibroblasts reside in different skin layers. Characterization of their electrotactic response for **i)** different EF magnitudes and **ii)** alternating EF directions is done in mono-culture (device μ1) and is compared to literature values. Co-culturing these cells requires adequate mixing and appropriate cell media to investigate if **(1)** these cells can be co-cultured and electrically stimulated and **(2)** if these cells show the same electrotactic behavior in mono- and co-culture.

For the first time, we demonstrate the concurrent electrotactic stimulation of co-cultured keratinocytes and fibroblasts. We present here two different microfluidic electrotaxis platforms to explore DC stimulation of human skin cells using salt-bridgeless electrodes in mono- and co-culture configurations. The first platform leverages soft lithography to yield consistent structures that individually enable electric field dosage dependency exploration of human keratinocytes and fibroblasts. This device allowed the initial determination of electrotactic thresholds as well as the influence of electrical polarity switching on cellular directionality dynamics in mono-cultures. The second platform exploits double-sided adhesives to permit straightforward co-culture seeding and cultivation while having a current divider design that provides six different EF intensities from a single pair of electrodes. This platform leverages larger supercapacitive electrodes to achieve DC stimulation for many hours and assess the concurrent dynamics of keratinocyte and fibroblast electrotaxis. The combination of both bioelectronic microfluidic platforms enables direct comparison of electrotactic metrics of both cell types when they are alone or coupled.

## 2. Results

### 2.1. Mono-culture electrotaxis on human skin cells

#### 2.1.1. Keratinocytes migrate toward the cathode

Both keratinocytes and fibroblasts were subjected to a step-wise increased EF magnitude to determine their threshold for electrotaxis. **Figure 2** depicts cell migration for keratinocytes and **Figure 3** for fibroblasts under the influence of an externally applied EF field within a single-channel microfluidic device (***μ1***). Manual cell tracking during time-lapse imaging provides insight into individual cell movement dynamics, showcased in hairline plots for cells in a non-stimulated state (control) and under an external stimulus (Figure 2A and Figure 3A). The individual cells are then analyzed collectively to determine their directedness in relation to the applied EF (**+1** being toward the **cathode, -1** toward the **anode**, c.f., section 5.8) and velocity. No adverse cellular reactions to the applied currents (i.e., cell death, impaired migration) were observed, indicating that electrical stimulation with a salt-bridgeless system is appropriate for the parameters and conditions applied here. For both cell types, the non-stimulated control showed random, non-directed movement, as is also expected in their normal state.

**Figure 2:**
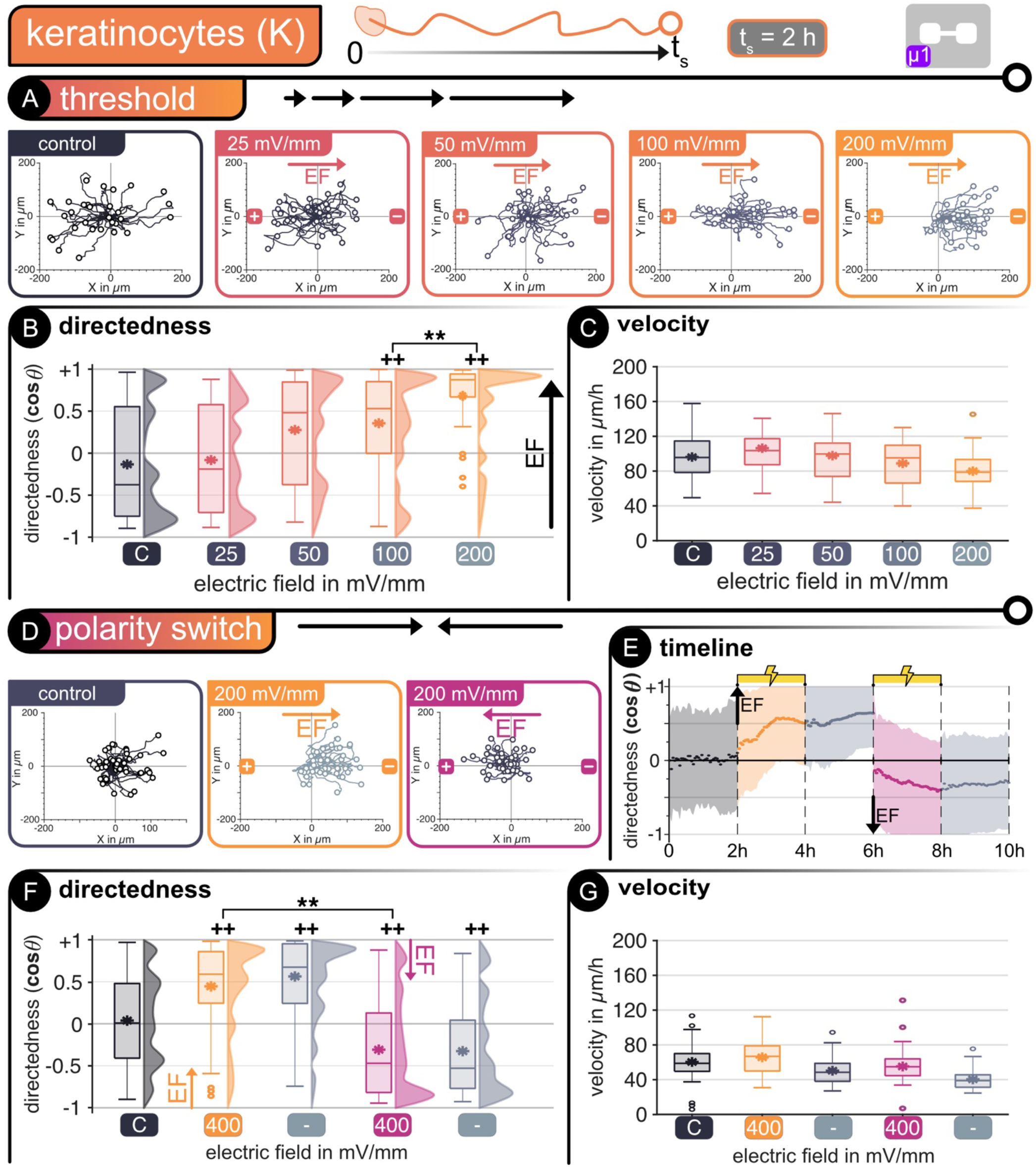
Keratinocytes - *Threshold determination*. **(A)** Cell tracking in microfluidic device *μ1*. Hairline plots represent individual movement over 2 hours for non-stimulated and stimulated cells at different electric fields between 25 – 200 mV mm^-1^ (n = 50). Migration statistics **(B)** directedness and **(C)** velocity. *Polarity switch* - **(D)** Hairline plots for movement over 2 hours for non-stimulated and stimulated cells at alternating electric fields of 200 mV mm^-1^ (n = 50). **(E)** Timeline of polarity switch experiment. Dots represent average every 3 min and shaded area represents standard deviation. Migration statistics **(F)** directedness and **(G)** velocity. | For all box plots, the box represents the 1st and 3rd quartiles, middle line shows the median, * denotes the mean value, the whiskers represent the max. and min. values, respectively, and circles represent outliers. ++ p < 0.01 when compared to the control with no EF. ** p < 0.01 when compared amongst stimulated cells of different EF.

**Figure 3:**
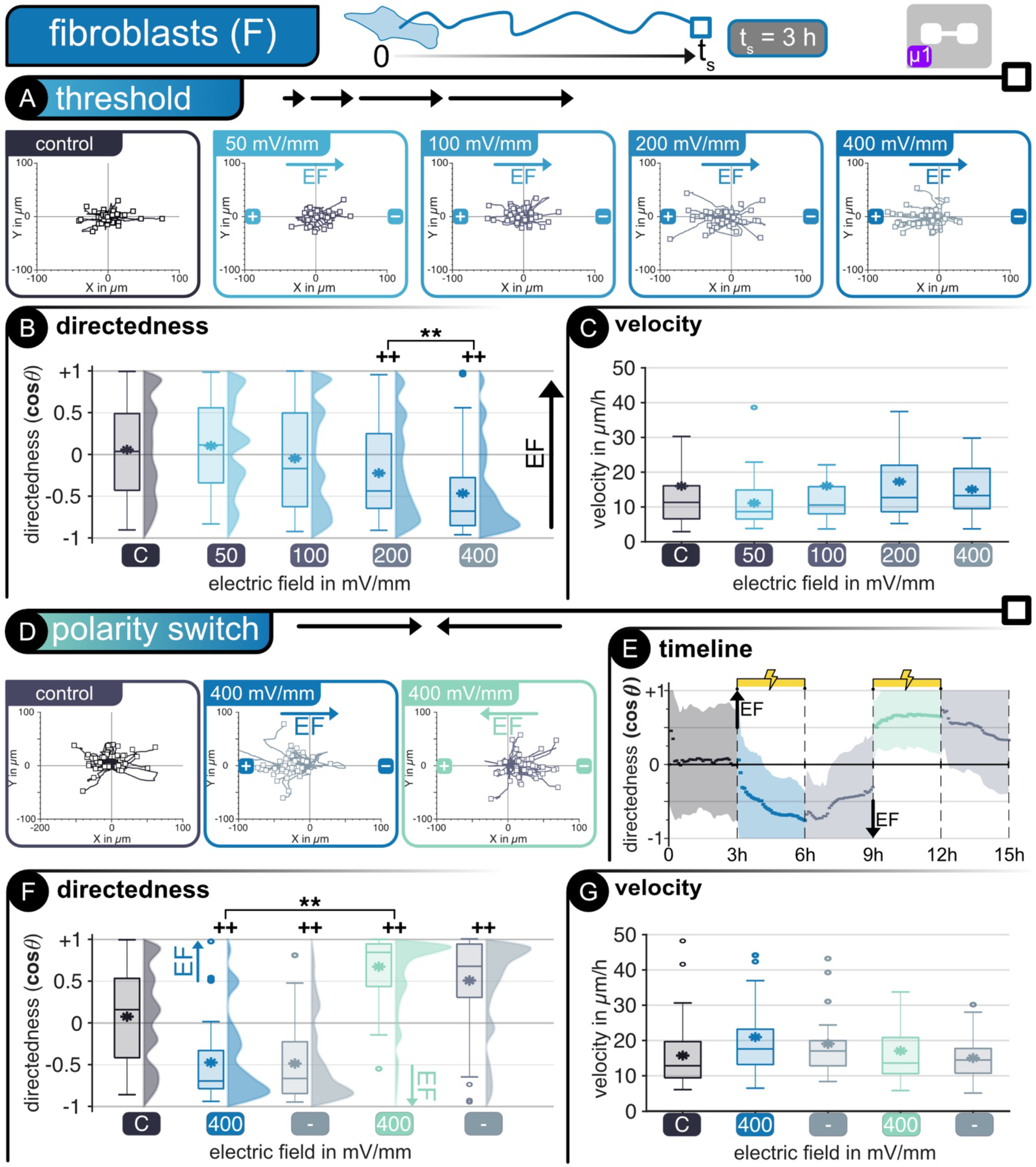
Fibroblasts - *Threshold determination*. **(A)** Cell tracking in microfluidic device *μ1*. Hairline plots represent individual movement over 3 hours for non-stimulated and stimulated cells at different electric fields between 50 – 400 mV mm^-1^ (n = 50). Migration statistics **(B)** directedness and **(C)** velocity. *Polarity switch* - **(D)** Hairline plots for movement over 3 hours for non-stimulated and stimulated cells at alternating electric fields of 400 mV mm^-1^ (n = 50). **(E)** Timeline of polarity switch experiment. Dots represent average every 3 min and shaded area represents standard deviation. Migration statistics **(F)** directedness and **(G)** velocity. | For all box plots, the box represents the 1st and 3rd quartiles, middle line shows the median, * denotes the mean value, the whiskers represent the max. and min. values, respectively, and circles represent outliers. ++ p < 0.01 when compared to the control with no EF. ** p < 0.01 when compared amongst stimulated cells of different EF.

Beginning at an EF of 100 mV mm^−1^, keratinocytes show directed migration toward the cathode during threshold determination reaching full directedness at an EF of 200 mV mm^−1^. This threshold is in accordance with previous reports (Figure 2B).^17,32^ There is a significant difference in directedness between the control cells (no stimulation) and those subjected to exogenous EFs. The differences in average directedness range from -0.21 for the control to +0.36 (p < 0.01) and +0.69 (p < 0.01) at 100 and 200 mV mm^−1^, respectively. There is also a significant difference between the distribution of directed cells between 100 and 200 mV mm^−1^. At lower EFs, the directedness of movement was not noticeably influenced, as seen in the hairline plots and average directedness for 25 and 50 mV mm^-1^, respectively. The velocity of keratinocytes is not significantly affected by the presence of an exogenous EF, such that all tested cells exhibit similar migration speeds of around 90 μm h^−1^ on average (Figure 2C).

After identifying the migration threshold for keratinocytes, they were subjected to alternating stimulation pulses at ±9.5 μA, thus 200 mV mm^-1^ with alternating directionality in device μ1. This experiment is intended to determine if the cellular movement could be switched at will within the microfluidic device using the same pair of electrodes. We could confirm that a switch in EF polarity also reversed the migration, with cells changing direction, migrating alongside the EF lines towards the cathode (Figure 2D). The reversed polarity of the applied stimulus had an evident impact on the directedness of the cells, with an initial random movement without stimulation followed by orientation towards the current cathode during the stimulation phases. Furthermore, after the first stimulation phase, residual directedness is noticeable during the following 2 hours when no stimulation is applied (Figure 2E). A significant difference exists between the random movement without stimulation and the directed migration with stimulation. During the initial stimulation phase (positive), the keratinocytes showed directed electrotaxis by changing their directedness from +0.05 to +0.49 (p < 0.01), with most cells moving towards the right side of the channel (cathode) (Figure 2F). After polarity reversal, most cells realigned towards the new cathode (left side). They directed their movement throughout the complete stimulation phase, with a new average directedness of -0.39, significantly differing from the non-stimulated cells (p < 0.01). During pauses between polarity switching, cells retained significant directedness compared to the non-stimulated control.

The average velocity for non-stimulated and stimulated keratinocytes was significantly lower during polarity switching experiments than monophasic stimulation (Figure 2G). This decrease could be due to the polarity-switching experiments performed at higher passages, leading to slower motility. ^33^ Nevertheless, the internal non-stimulated control still justifies the results showing evident directional switching migration of keratinocytes. The results obtained herein are equivalent to previously published investigations on human keratinocytes, thus ensuring that our salt-bridgeless system does not negatively influence the electrotaxis of these cells.^17^

#### 2.1.2. Fibroblasts migrate toward the anode

Fibroblasts are larger and less mobile than keratinocytes, as evidenced by the shorter distance traveled by these cells (Figure 3A). Furthermore, fibroblasts require higher EFs and longer stimulation times to show directed electrotaxis. Starting at 200 mV mm^−1^, directed migration toward the anode is noticeable compared to the non-stimulated cells, with changes in average directedness from +0.05 to -0.31 (p < 0.01). Increasing the stimulation EF to 400 mV mm^−1^ leads to a more defined distribution of directed cells with a more pronounced directedness of -0.52, which significantly defers from the non-stimulated (control) and the 200 mV mm^−1^ field (p < 0.01). The directed movement influenced by electrical stimulation was not noticeable below 200 mV mm^-1^ with directedness values similar to the non-stimulated cells of +0.06 and -0.08 for 50 and 100 mV mm^-1^, respectively (Figure 3B). On average, all fibroblasts migrate at approximately 15 μm h^−1^ without any significant difference between non-stimulated and stimulated cells (Figure 3C).

The influence of polarity reversal during direct current stimulation on fibroblasts was studied at higher currents of ±15 μA, producing EF values of 400 mV mm^-1^ in alternating directions within device μ1. These cells, analogous to keratinocytes, retain their previously observed electrotactical properties with a clear orientation towards the anode when an external stimulus is applied (Figure 3D). During the initial phase without stimulation, fibroblasts move randomly, as expected, changing their directedness opposite to the EF lines as soon as stimulation is applied. During the first stimulation phase (positive EF), fibroblasts migrated toward the anode (left). Reversal of the polarity leads to a subsequent reversal in electrotaxis, with most cells reorienting themselves toward the new anode (right). The average directedness of fibroblasts increased with sustained stimulation. This observation is valid for both phases with stimulation. Like keratinocytes, the fibroblasts retain their directedness long after the stimulation is interrupted and slowly transition back to more random movement, similar to their non-stimulated behavior (Figure 3E). The differences in cellular directedness were significant when compared to non-stimulated cells, with changes in average values from +0.11 (control) to -0.65 and +0.72 for the positive and negative stimulation phases, respectively. Significant changes in directedness compared to the control were also noticeable during the pauses after stimulation phases (Figure 3F). An average migration velocity of 17 μm h^-1^ was determined for fibroblasts with slightly higher velocity during stimulation (Figure 3G). These results confirm previous observations on human fibroblasts.^6^

Both tests on keratinocytes and fibroblasts demonstrate that a salt-bridgeless system supplying direct current stimulation to the cell media can elicit the expected electrotactical response for both epithelial cells. Furthermore, these results show no noticeable adverse effects on cell mobility or viability. These results provide the baseline for a direct comparison between mono- and co-cultured keratinocytes and fibroblasts.

### 2.2. Co-culture electrotaxis of human skin cells

#### 2.2.1. Finding a suitable co-culture media

When establishing a co-culture, it is essential to consider a suitable media that accommodates different cell types. In this study, we focused on the cell media previously employed in mono-cultures to maintain the chemical composition and, thus, the electrical properties of the media the same. The suitability of the media was assessed through imaging (**Figure 4A**) and metabolic assessment (Figure 4B) (c.f., section 5.7), as cells need to show similar metabolism, morphology, and phenotypes to ensure comparability between mono- and co-culture stimulation. Three media were investigated for mono- and co-cultures of keratinocytes (*K*) and fibroblasts (*F*), mainly serum-free K media (M1), serum-containing F media (M2), and a 1:1 mixture of both (M3).

**Figure 4:**
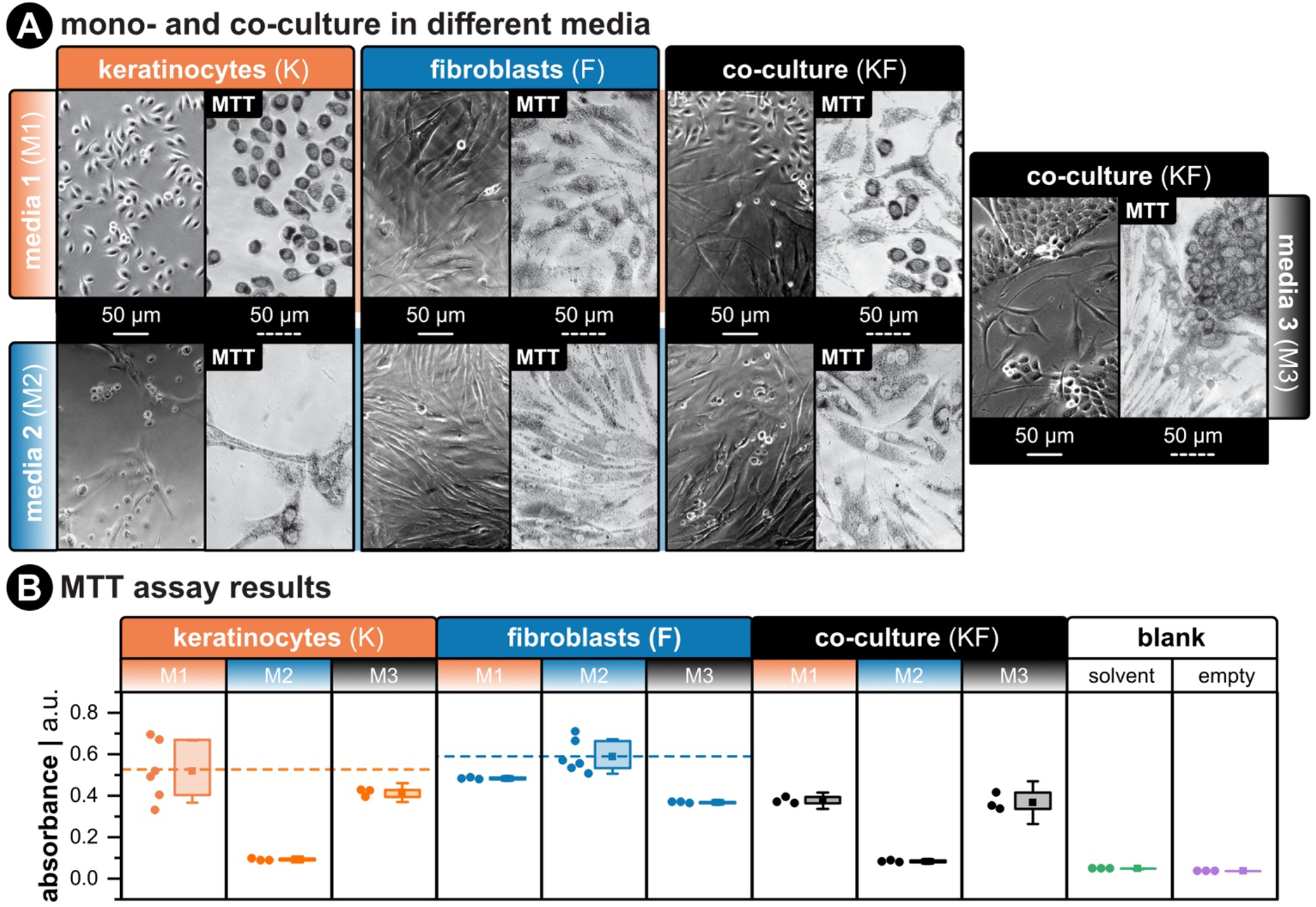
**(A)** Phase-contrast images of keratinocytes, fibroblasts, and a mix of both types in different media. Left panel shows cultured cells 36 h post-seeding. Right panel shows cells with reduced formazan crystals before placing on the plate reader. All cells were seeded into 96-well plates and at the same total number of cells (1 × 10^5^) with the same volume (100 μL). For co-culture wells, the total amount was the same (5 × 10^4^ for K and F). All scale bars are 50 μm. **(B)** Metabolic activity measured by absorbance at 570 nm. More signal correlates to more metabolic activity. The horizontal dashed line for each group corresponds to mean absorbance for the control group (i.e., the media used during mono-culture experiments). The “blank” group shows absorbance results for wells filled with the MTT solvent (i.e., dimethyl sulfoxide - DMSO) or wells that were left empty.

We found that *Ks* were more sensitive to media composition than *Fs* and required keratinocyte media (M1) to seed and proliferate effectively. *Ks* in fibroblast media (M2) retained a rounded morphology and did not show the characteristic extension of lamellipodia on the substrate. On the other hand, *Fs* could seed and proliferate in all media but showed less metabolic activity in M1 and M3 compared to control media M2. Co-culture (*KF*) results showed that *Ks* required supplements in M1 to seed and be metabolically active, while mixed media (M3) and M1 were adequate for co-culture metabolic activity. In keratinocyte media (M1), both cell types formed an evenly mixed population, but *Fs* did not have the structured alignment they possess in fibroblast media (M2). In mixed media (M3), *Fs* regained their structured phenotype, and *Ks* formed more cell-cell junctions.

#### 2.2.2 Directional separation of keratinocytes and fibroblasts

After carefully evaluating cell viability and metabolism in an adequate medium for keratinocytes (*Ks*) and fibroblasts (*Fs*) in co-culture, these cells were investigated jointly in a common bioelectronic platform. The co-culture was seeded in a multi-channel microfluidic device (**μ2**) with branching channels through which current at different magnitudes could flow, thus generating a variety of carefully defined EFs to guide cellular migration. Using a drop-cast seeding method of both cell types simultaneously in device μ2 led to the homogeneous distribution of *Ks* and *Fs* within all device channels in all biological replicates, thus allowing reproducible experimentation. The morphology of the cells in the co-culture within the microfluidic channel was similar to the preliminary observations during MTT assays in well plates (Figure 4A). Fibroblasts were independent, as expected from *F* mono-cultures, while keratinocytes were evenly spread between independent cells and *K* clusters. This difference between mono- and co-cultured *Ks* did not impact their response to the applied EF. All stimulated cells were reactive to the EF guidance cues throughout the 12 h stimulation.

*Ks* and *Fs* demonstrated similar behavior in co-culture as in mono-culture, with a characteristic anodic migration for *Ks* and cathodic migration of *Fs*, effectively separating both cell types when stimulation was applied in co-culture (**Figure 5**). The presence of both cells within the microfluidic channel had no evident influence on their electrotactic properties. In some situations, migration was slightly hindered through cell-cell collisions, but overall directedness and velocity were unaffected. Though cell density in each channel varied slightly, the influence of the applied EF was clearly distinguishable between *Ks* and *Fs*. Simultaneous cell tracking and overlapping of the hairline plots emphasizes differences amongst these cells in the direction of migration, their migration path, and their velocity (Figure 5A). The differences in cell size and mobility are further noticeable during imaging, with clearly spread out *Fs* slowly moving through the microchannels and both individual and agglomerated *K*s swiftly migrating and reorienting themselves. (Refer to supplementary videos for individual EFs)

**Figure 5:**
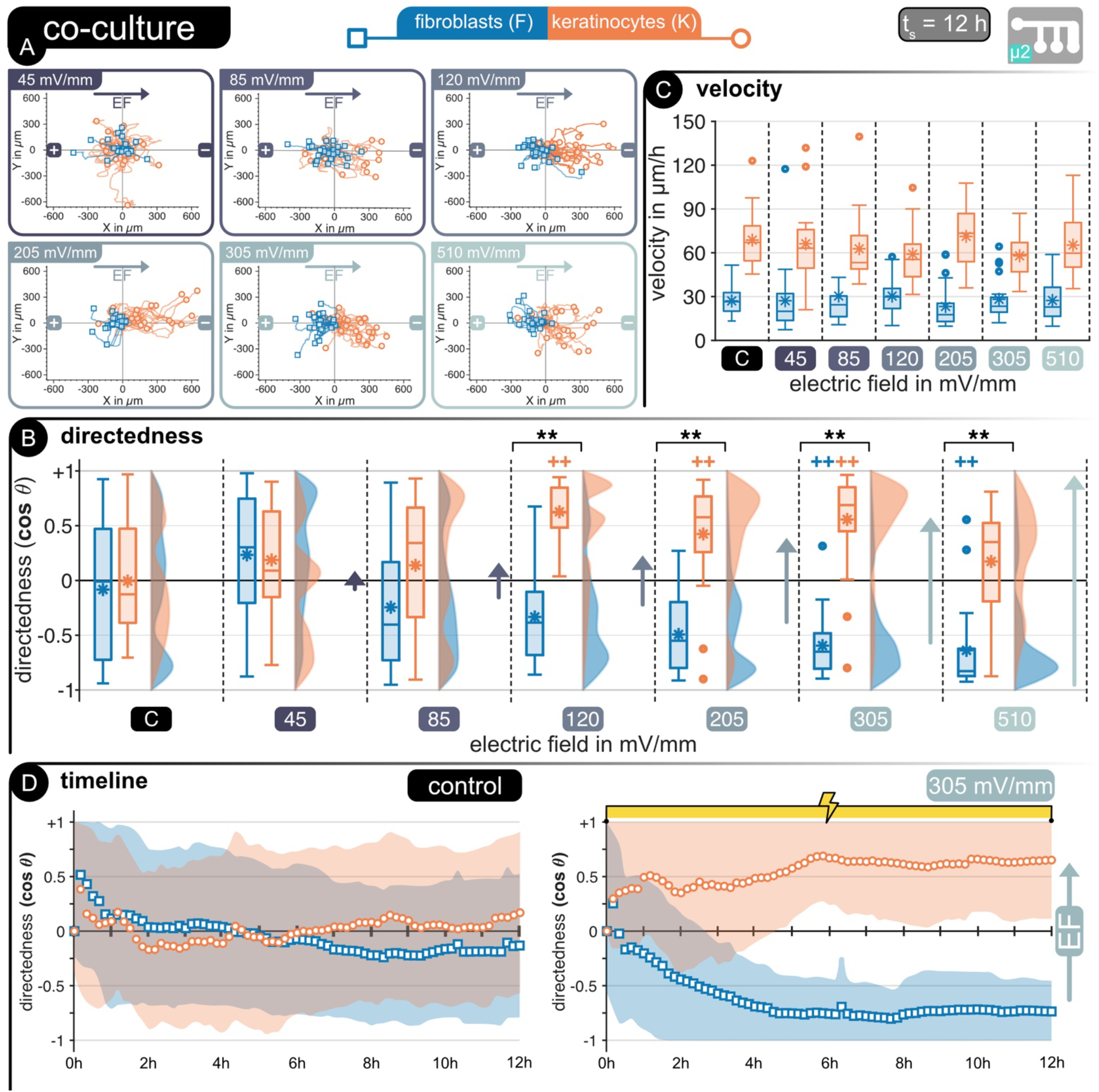
**(A)** Cell tracking in microfluidic device *μ2* for keratinocytes (orange) and fibroblasts (blue). Hairline plots represent cell movement over 12 hours of stimulated cells at different electric fields between 45 – 520 mV mm^-1^ (n = 25 / cell type). Migration statistics **(B)** directedness and **(C)** velocity for keratinocytes (orange) and fibroblasts (blue). **(D)** | For all box plots, the box represents the 1st and 3rd quartiles, middle line shows the median, * denotes the mean value, the whiskers represent the max. and min. values, respectively, and circles represent outliers. ++ p < 0.01 when compared to the control with no EF. ** p < 0.01 when compared amongst cell types at same EF strength.

Analogous to the mono-cultures (Figures 2 and 3), a clear impact of the EF magnitude on the directedness of the cells was notable in co-culture, with *Ks* showing significant directedness in comparison to non-stimulated cells at lower EFs than *Fs*. Both cell types showed similar thresholds in co-culture as in mono-culture, with a significant difference from the control group starting at 120 mV mm^-1^ for *Ks* and 305 mV mm^-1^ for *Fs* (Figure 5B). Co-cultures were stimulated for more extended periods than mono-cultures leading to more pronounced directedness at EFs below the previously established thresholds, similar to previous observations in the literature about mono-cultures.^17,34^ Increasing the EF magnitude led to more defined directedness in *Fs* analogous to observations in mono-cultures. However, this effect was not observable for *Ks*, with cells losing directed migration at the highest EF applied.

*Fs* and *Ks* showed no significant change in their migration velocity under the influence of an externally applied EF compared to control cells, as expected from the individual investigation of each cell type. However, a difference in velocity between the mono-cultured and the co-cultured cells is notable, particularly for Fibroblasts. These cells increased their migration velocity by approximately 50% to 23 μm h^-1^ when in co-culture, whereas the Keratinocytes retained similar speeds as the mono-cultured ones. (Figure 5C).

A further distinction between the co-cultured *Fs* and *Ks* and their mono-cultured counterparts is noticeable in the changes of directedness over time, particularly for higher EFs, where both cells show significant differences from non-stimulated cells. Figure 5D shows a comparison between *Fs* and *Ks* directedness without stimulation (control) and under an EF of 305 mV mm^-1^. Non-stimulated cells show random movement for 12 hours, while stimulated cells slowly direct their movement alongside (*Ks*) and opposite (*Fs*) to the EF lines. This alignment is slower for co-cultured cells than for mono-cultured ones reaching a maximum directedness after 6 h of stimulation. Nonetheless, a defined cellular separation is notable, as expected from the individual behavior of each cell type. These results respond both questions of this work (Figure 1), as *Fs* and *Ks* could be successfully cultured together, retaining their electrotactic properties and aligning themselves to externally applied EFs. Furthermore, both cell types showed an analogous electrotactic response in co-culture as in mono-culture.

## 3. Discussion

Human keratinocytes (*Ks*) and fibroblasts (*Fs*) reside in different layers of the human skin and are, therefore, never found in planar co-cultures. Nonetheless, it has been shown that these cell types concurrently influence each other during the healing process of wounds, and the underlying mechanisms still need to be elucidated. ^35,36^ The electrotactic behavior of human fibroblasts and keratinocytes, as well as the suspected underlying mechanisms that lead to the directed migration alongside an EF, such as calcium channel activation and PI3K signaling, have been previously proposed and investigated.^6,37^ However, these experiments have usually been done on one cell type at a time, thus relinquishing the active interactions between different cell types in their usual biological environments. Limitations here have been mainly due to difficulties of concurrent growth of different cell types within microfluidic devices and the employed stimulation setup. In this study, the electrotactic behavior of both cell types was carefully investigated first individually, which serves as a starting point to determine the adequacy of a salt-bridgeless stimulation setup to study the electrotactic response of human skin cells. Second, their electrotactic behavior was reproduced and evaluated in co-culture experiments.

Co-cultured fibroblasts and keratinocytes were successfully stimulated within device μ2 and reacted as expected during direct current stimulation, with clear distinction in the directedness of movement for both cell types. Furthermore, both cells began electrotaxis at the expected EF strengths, as determined during the initial threshold experiments. One of the key points we want to highlight in this work is that neither the seeding method, the stimulation protocol, nor the employed salt-bridgeless electrodes harmed the cells but rather confirmed and expanded on electrotaxis dose-response findings of human keratinocytes and fibroblasts. Therefore, a clear cellular partition was achievable under the chosen biological and electrical conditions (Figure 5). Keratinocytes migrated distinguishably towards the cathode, regardless of the new morphological phenotype (clusters) obtained in the new media, while fibroblasts concurrently migrated in the opposite direction towards the anode. The observed behavior is not representative of how wound healing typically happens, with an initial keratinocyte migration towards the wound center for re-epithelization, followed by the underlying fibroblast migration for restructuring. However, these newly validated platforms can be further leveraged to investigate how different biological factors are orchestrated (e.g., plasma, serum transition) during typical wound healing mechanisms.

Experimentation of these cell lines in co-culture under the same stimulation paradigms has not been demonstrated before.^38^ As these cells require different conditions to be viable in culture, identifying a suitable cell media for coexistence *in vitro* was fundamental. Here three main factors were considered: i) the new media needed similar calcium concentration as the media for individual cells; ii) it should allow for clearly distinct morphological phenotypes between cell types (if no fluorescent tagging or cell-specific surface reporters are used), thus allowing differentiation amongst cell types during tracking; iii) and it should not significantly diminish the metabolic activity of the cells. After several combinations, a one-to-one ratio between fibroblast and keratinocyte media was chosen. Both cell types were metabolically active, had comparable calcium concentrations required for electrotaxis, and could be distinguished from each other. Fibroblasts showed similar morphology as in their individual media, while keratinocytes formed two phenotypes of individual keratinocytes and keratinocyte clusters (KCs) (Figure 4).

This higher degree of KCs directly correlates to the media’s calcium concentration.^39^ In mammalian skin, there is a calcium gradient from the apical layer (highest calcium concentration with terminally differentiated keratinocytes and strong cell-cell adhesion) to the basal layer (lowest calcium concentration with proliferating keratinocytes and weak cell-cell adhesion).^40,41^ Also, fibroblasts require extracellular calcium concentrations above 1.4 mM to proliferate.^40^ In culture, the same modulation of cell-cell adhesions can be adjusted with calcium concentration.^37,42^ Note that the calcium chloride (CaCl_2_) in keratinocyte media (M1), which is a basal media, is 0.06 mM, and in fibroblast media (M2), it is 1.80 mM. Therefore, in the mixed media (M3), it was set to 0.93 mM. The compromise of extracellular calcium concentration on keratinocyte and fibroblast proliferation explains why for the same number of cells, the metabolic activity was lower in the co-culture with M3 compared to *Ks* in M1 and *Fs* in M2. The combination of the metabolic activity results and the formation of distinct subpopulations of *Fs* and *Ks* with strong cell-cell adhesions, which favors cell tracking, led us to choose the mixed media (M3) for all co-culture experiments.

The results obtained during the initial experiments with device μ1 are on par with previously postulated electrotactic parameters for both *Ks* and *Fs*.^16,17,34,43^ These results further emphasize that a salt-bridgeless system does not negatively influence cellular behavior, facilitating the experimental setup. The fundamental characterization of keratinocytes (Figure 2) and fibroblasts (Figure 3) provides valuable insights for the future development of bioelectric wound dressings aimed at wound healing acceleration through cell migration redirection. When stimulating *Ks* and *Fs* separately, the experiment can continually be optimized to the respective cell type. We argue that co-culture experiments have an essential role to fill, as these results demonstrate that a relevant response can be triggered in both types of cells, even when trade-offs have to be made between what is optimal to impact *Ks* and *Fs*, respectively. It is a small but crucial step closer to real-world application. It should be noted that the differences in migration direction, cathodic for keratinocytes and anodic for fibroblasts, are known behaviors for which a clear explanation is still missing. Several hypotheses have been postulated trying to explain the directional choice of these cells involving calcium signaling pathways,^40,41,44^ PI3 kinase (PI3K),^6,45^ transforming growth factor-β3 (TFG-β3),^46^ Golgi polarization,^47^ or integrin expression,^48^ nonetheless, a clear.

Skin wounds might seem inconsequential in everyday life for most people, as the skin heals itself over days to weeks without much conscious maintenance. For the skin itself, however, this process requires a plethora of different cell types, signaling molecules, and communication pathways as it undergoes the four stages of hemostasis, inflammation, proliferation, and remodeling.^49^ A better understanding of the underlying cellular mechanisms involved in electrotaxis, as well as technical solutions that allow translation from *in vitro* testing to *in vivo* applications, are required to develop successful clinically validated wound therapy based on direct current stimulation.^50–59^ This study discusses two devices and electrodes designed to facilitate electrotaxis experimentation. Both devices have advantages and drawbacks and can be leveraged based on the desired purpose. The PDMS and PMMA devices used in the study have proven reliable for electrotaxis research. They provide a suitable environment for cell seeding and proliferation and enable direct current stimulation of different cell populations without the need for salt bridges. The second device and its drop-cast seeding approach make co-culture seeding and experimentation straightforward, allowing for future exploration of more biologically relevant constructs.

We believe this work is just the beginning of studying co-cultured keratinocytes/fibroblasts electrotaxis, as the technology devised herein can significantly facilitate further experimentation in co-culturing these cells under different biological conditions in order to identify which limits and cues govern cellular migration in real wounds. Now the sandbox is filled for other researchers to play and explore the bioelectric communication and mechanisms between these two cell types.

## 4. Conclusion

In this work, we demonstrate that human fibroblasts and keratinocytes can be electrotactically guided in a salt-bridgeless system concurrently, retaining their expected electrotaxis, thus leading to the directional separation of both cell types. We leveraged two different microfluidic systems compatible with different salt-bridgeless electrodes that facilitate *in vitro* experimentation with cell cultures, particularly on the effects of DC stimulation on cellular electrotaxis. All cells tested in this study showed the expected electrotactic behavior under the influence of an externally applied EF with directedness towards the cathode for keratinocytes and the anode for fibroblasts regardless of if they were in mono- or co-culture. Furthermore, both cells could be guided at different EF strengths and polarities. A suitable media combination was determined for the viability of the cells in co-culture and was found to lead to an evident change in keratinocyte agglomeration. This study demonstrated for the first time the concurrent seeding, growth, and electrotactic stimulation of the co-culture of these cells in a microfluidic device. We propose a new approach for both cell seeding and direct current stimulation for the concurrent investigation of both cell types involved in several stages of wound healing, thus opening the door to faster characterization and future clinical *in vivo* applications of direct current stimulation therapy.

## 5. Methods

### 5.1. Compact microfluidic platforms with salt-bridgeless electrodes

Microfluidic devices allow precise control of the electric field (EF) distribution through accurate fabrication, known media composition, and precise current control, thus ensuring that all cells studied are subjected to the same stimulus. This is possible as the EF inside a rectangular channel, as the ones employed in this study, is directly dependent on the channel’s cross-section (width - *w*, height - *h*), the conductivity of the fluid in the channel (*σ*), and the applied current *I* following Ohm’s law:

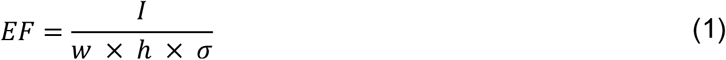

Here we utilize two approaches to microchannels in the form of a single-channel (**μ1**) and a multi-channel (**μ2**) device (**Figure 6A**). The determination of the applied EF is straightforward in device μ1, as the cross-section is well-defined and current flows through the channel, the EF distributes homogeneously across the experimental area. The small cross-section achievable through soft-lithography on PDMS leads to high EFs utilizing a relatively small current. However, the current must be sequentially changed to achieve different EFs with this device. On the other hand, the multi-channel microfluidic device μ2 provides six different EFs utilizing one set current. It serves as a current divider and subsequently an EF divider, resulting in 6 different regions with EF ratios of 13 : 8 : 5 : 3 : 2 : 1, as validated through FEA analysis (Figure 6B).

**Figure 6:**
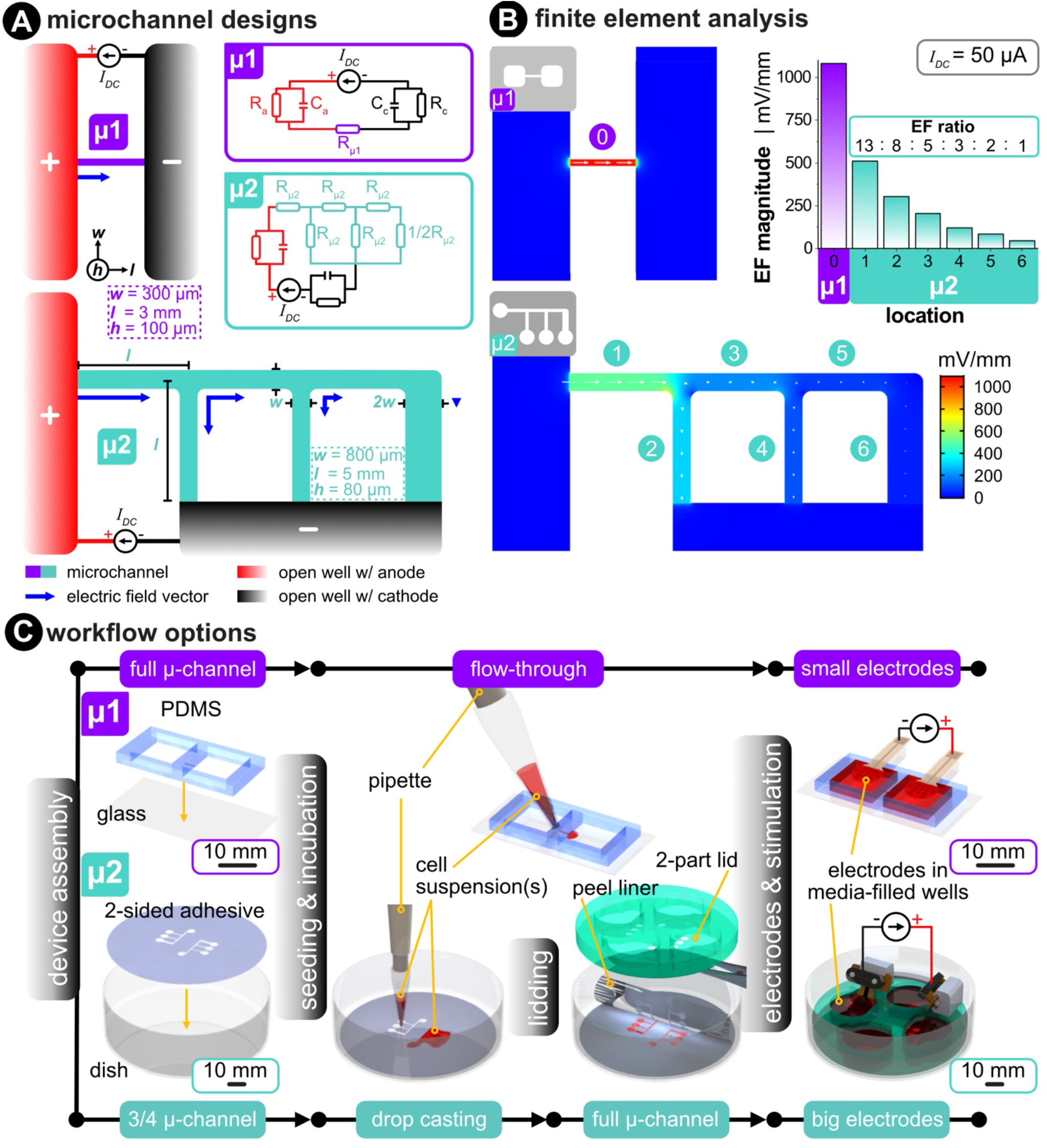
**(A)** Device **μ1** (violet) shows a top view of a simple rectangular microchannel that connects two open wells, each filled with excess media (mL in the wells and μL in the microchannel). Device **μ2** (turquoise) shows a branching microchannel network that forms a current divider. Insets of the electrical equivalent circuit relate the microchannels to resistors and include the electrode-electrolyte interface of both the anode and cathode. (*R* = resistance, *C* = capacitance, and *I*_*DC*_ = DC source). **(B)** Three-dimensional finite element analysis of both microchannel designs. The same input current (*I*_*DC*_ = 50 μA) was used for both cases. The color scale and white arrows signifies the EF magnitude and direction, respectively. **(C)** Two different workflow options used in this paper allow for protocol flexibility. μ1 is made via soft lithography of molded polydimethylsiloxane (PDMS) that is irreversibly bonded to glass via air plasma exposure. μ2 is made by bonding the bottom side of a laser-structured double-sided adhesive to a Petri dish. Cell suspension(s) are flow-through seeded in μ1 and drop-casted in μ2. After cell seeding and growth in μ2, the adhesive’s top protective liner is removed, and an acrylic lid is added to complete the microchannels. Once the desired cell confluency is obtained, electrodes are added and stimulated in an incubated microscope.

### 5.2. Fabrication of microfluidic devices

#### 5.2.1. Device 1: more complex but higher geometrical precision

These devices were fabricated according to protocols according to Leal et al., as a guide, please see Figure 6C.^29^ The microfluidic channels (3 × 0.3 × 0.1 mm^3^ – l × w × h) were fabricated through soft lithography of two-component polydimethylsiloxane (PDMS) Sylgard 184 (Dow Corning, MI, USA) onto SU-8 structures. The PDMS was heat cured at 65°C, and subsequently, the independent structures and the media reservoirs (10 × 10 × 2 mm^3^) were cut. Finally, the devices were plasma treated for 2 minutes at 300 W, air flow of 10 sccm, and a pressure of 0.8 mbar before being irreversibly bonded to a glass slide. The addition of OH groups on the PDMS walls increases the hydrophilicity of the channel, thus facilitating flow-through cell seeding.

#### 5.2.2. Device 2: more flexibility and larger feature size

As a guide, please see Figure 6C. Besides the sterile polystyrene dish, all microfluidic device components are fabricated with a 30 W carbon dioxide (CO_2_) laser (Universal Laser Systems, VLS 2.30). For the acrylic-based double-sided pressure-sensitive adhesive (Adhesives Research, 90445Q), a kiss-cut was made with 7.5 W at 70 mm s^−1,^ and a through-all cut was made with 24 W at 70 mm s^−1^. The bottom-side liner (i.e., without the kiss-cut) was first peeled off to expose the bottom-side adhesive, then was pressure-bonded by hand to a new Petri dish. Importantly, batches of dish/adhesive were placed in a vacuum desiccator overnight to remove any air bubbles during bonding. These are stored on the shelf until further use. The acrylic (Modulor, Germany) two-part lid consists of a thin 0.5 mm base that only has fluidic vias and a thicker 8.0 mm reservoir-defining layer (Figure 1c, pink part). These two parts were solvent-bonded together using dichloromethane (Modulor, Germany). Note that this lid is not bonded to the microfluidic adhesive until the cells are seeded (see section 5.6).

### 5.3. Seeding of microfluidic devices

#### 5.3.1. Device μ1: flow-through seeding

The protocol used here follows Leal et al.^29^ The PDMS devices were immersed in 70 % ethanol for 30 minutes for sterilization and then placed in sterile PBS (Carl Roth GmbH + Co. KG, Karlsruhe, Germany) for 1 hour before cell seeding. Before the seeding procedure, the devices were dried, and any remaining PBS inside the channel was carefully aspirated. Subsequently, a 10 μL droplet with approximately 10^3^ cells μL^−1^ was pipetted at the entrance of the microchannel, and the media and cells were driven within the channel through capillary forces. The devices were subsequently placed in an incubator (37°C, 5 % CO_2_) for 3 h to permit cell attachment on the glass. After cell adherence, the media reservoirs were filled with 200 μL of the corresponding media for each cell line, M1 for keratinocytes and M2 for fibroblasts (see section 5.4).

#### 5.3.2. Device μ2: drop-cast seeding

The protocol used here follows Shaner et al.^22^ Petri dish devices were first washed with 70 % ethanol and nitrogen dried. Then the devices were air plasma-treated (Femto, Diener Electronics) on the same day as seeding to improve cell adhesion. The settings were 30 W, 3 min, and 10 sccm air. The dishes were soaked in 70 % practical grade (p.a.) ethanol under the cell culture hood (Safe 2020, Thermo Scientific) for 15 min before washing with sterile water, then allowed to dry in the hood. Lastly, the devices were subjected to the hood’s integrated UV lamps for 1 h. Sub-cultured keratinocytes or fibroblasts were harvested into the desired concentration (5 × 10^5^ cells mL^−1^). A volume of 100 μL of this suspension was seeded (5 × 10^4^ total cells) directly onto the open microchannels and incubated for 3 h to allow for cell attachment. Afterward, 10 mL of fresh media is added and placed back into the incubator for one to three days. After the desired confluency was achieved, the media was aspirated until only a small amount of media resided in the microchannels leaving the liner as dry as possible. The liner was then peeled, and the two-part acrylic lid was aligned and fixed using alignment marks etched into the adhesive. Media was immediately replenished by initially flowing 100 μL directly into the microchannels to displace trapped air, and then the wells were filled with more fresh media. The electrodes were assembled and placed into the reservoir, and the corresponding wires were routed through the lid, which was applied to prevent evaporation

### 5.4. Finite Element Analysis (FEA)

The microfluidic devices were designed and exported (IGS file extension) in Solidworks (version 2021). COMSOL Multiphysics^®^ software (version 5.3) was used to simulate EF distribution using the Electric Currents module. For EF distribution, electrodes sat on top of the reservoirs and were modeled to have the electrical conductivity of PEDOT:PSS hydrogels (*σ* = 2000 S m^−1^).^60^ The media was modeled with an electrical conductivity (*σ*) of 1.5 S m^−1^, which was measured for all three media options using a portable conductivity meter (DiST6 EC/TDS, Hanna Instruments, Germany). The relative permittivity of the media was 80, which is typical for saline water. The cathode was set to 0 V. The input current density (placed at the face of the anode) was swept to identify which input current is needed to achieve the desired EF strengths.

### 5.5. Fabrication of electrodes

#### 5.5.1. Metal electrodes coated with conducting polymer

The protocol used here follows Leal et al.^29^ In short, thin-film electrodes consisting of 300 nm platinum tracks with 700 nm of Sputtered Iridium Oxide Film (SIROF) active sites sandwiched between 10 μm of polyimide insulating film were fabricated in a Class 3 clean room at the University of Freiburg. These devices had an active electrode area of 20 cm^2^ onto which the conducting polymer poly(3,4-ethylenedioxythiophene) polystyrene sulfonate (PEDOT/PSS) was electrochemically polymerized. This was done from an aqueous solution containing sodium polystyrene sulfonate (NaPSS, 5 mg mL−1) and 3,4-ethylenedioxythiophene monomers (EDOT, 0.01 M) (Sigma Aldrich, MO, USA). The electropolymerization was done with a high-precision potentiostat/galvanostat (PGSTAT204, Metrohm Autolab B.V., Filderstadt, Germany). A three-electrode setup was employed in which the probe to be coated served as the working electrode (WE), a silver/silver-chloride (Ag/AgCl, BASI, USA) electrode as the reference (RE), and a stainless-steel sheet (≈2 cm^2^) as the counter electrode CE. The WE was driven at 0.9 V while the charge passing through the electrode was measured and utilized as a proxy to determine the polymer thickness. All electrodes were coated until 60 μC was reached, equivalent to a charge density of 300 mC cm^−2^.

#### 5.5.2. Non-metal electrodes coated with conducting hydrogel

The protocol used here follows Shaner et al.^30^ In short, the base electrode material was fabricated on the surface of thin sheets (75 μm) of polyimide (Kapton HN, Dupont, USA) and was carbonized using a mid-IR (wavelength of 10.6 μm) CO_2_ laser (VLS 2.30, Universal Laser Systems, USA). This process yields a material called laser-induced graphene (LIG). The LIG was coated with a pure PEDOT:PSS hydrogel to improve electrochemical properties. Specifically, the PEDOT:PSS dispersion (1.3 % in water) was spiked with 15 % dimethyl sulfoxide (DMSO) and cast onto the amine-functionalized and polyurethane-coated LIG, which improves adhesion between the LIG and hydrogel. PEDOT:PSS hydrogel-coated LIG electrodes were stored in 1× phosphate-buffered saline (PBS) until further use.

### 5.6. Cell culture media

Human epidermal keratinocytes immortalized with HPV-16 E6/E7 were courtesy of Prof. Dr. rer. nat. Thorsten Steinberg (Department of Dental, Oral and Jaw Medicine; University Clinic Freiburg). Keratinocytes were cultured in serum-free keratinocytes growth medium (KGM2, PromoCell, #C-39016) supplemented with bovine pituitary extract, epidermal growth factor, insulin, hydrocortisone, epinephrine, transferrin, and CaCl_2_ provided by the same manufacturer (SupplementMix, PromoCell, #C-20011), as well as neomycin (Sigma-Aldrich, #N1142) at final concentration 20 μg mL^−1^ and kanamycin (Sigma-Aldrich, #K0254) at final concentration 100 μg mL^−1^. This keratinocyte media is referred to as M1 in Figure 4. Human primary fibroblasts were courtesy of Prof. Dr. rer. nat. Thorsten Steinberg (Department of Dental, Oral and Jaw Medicine; University Clinic Freiburg). Fibroblasts were cultured in low-glucose Dulbecco’s modified Eagle’s media (DMEM, Sigma-Aldrich, #C-22320022) with 10 % fetal bovine serum (FBS, Sigma-Aldrich, # F0804). This media was supplemented with the same concentration of neomycin and kanamycin as M1. This fibroblast media is referred to as M2 in Figure 4. The final media used was a 1:1 mixture of M1 and M2. This co-culture media is referred to as M3 in Figure 4. Cell culture was incubated at 37°C and 5 % CO_2_ at 95 % humidity and routinely passaged when 80 to 90 % confluency was reached. Growth medium was exchanged three times per week. All experiments in this work included keratinocytes between passages 14 – 40 (low passages 14 – 33 were used for threshold characterization, and high passages 34 – 40 were used for other experiments) and fibroblasts between passages 9 – 11.

**Table 1:** Composition of media used in mono- and co-culture experiments and MTT assay.

| Code Name<br>(Fig. 4) | M1 | M2 | M3 |
| --- | --- | --- | --- |
| Abbreviation | KGM2 | DMEM | 50% KGM2 + 50% DMEM |
| Additives | - bovine pituitary extract<br>- epidermal growth factor<br>- insulin<br>- hydrocortisone<br>- epinephrine<br>- transferrin<br>- CaCl<sub>2</sub><br>- neomycin (20 $\mu\text{g mL}^{-1}$)<br>- kanamycin (100 $\mu\text{g mL}^{-1}$) | + 10% fetal bovine serum<br>+ neomycin (20 $\mu\text{g mL}^{-1}$)<br>+ kanamycin (100 $\mu\text{g mL}^{-1}$) | (50%)<br>- bovine pituitary extract<br>- epidermal growth factor<br>- insulin<br>- hydrocortisone<br>- epinephrine<br>- transferrin<br>- CaCl<sub>2</sub><br>(50%)<br>+ 10% fetal bovine serum<br>(100%)<br>± neomycin (20 $\mu\text{g mL}^{-1}$)<br>± kanamycin (100 $\mu\text{g mL}^{-1}$) |
| Cell Type | Keratinocytes | Fibroblasts | Keratinocytes + Fibroblasts |

### 5.7. Metabolic activity assay

Co-culture media options were evaluated via a cellular metabolic activity assay (i.e., MTT assay). First, cells were seeded (10^6^ cells mL^−1^ in 100 μL) in a 96-well plate. The plate was incubated for two days; then, the MTT assay was performed. The MTT reagent (mono-tetrazolium salt) was dissolved in sterile 1× PBS (5 mg mL^−1^. This solution was filtered through a 0.2 μm filter via a syringe into a sterile container. 10 μL of the MTT solution was added to the 100 μL-filled wells, then incubated for 3 h at 37°C. Since the media contains phenol red, the media was carefully aspirated, and the salt crystals were dissolved with 100 μL of sterile 100 % DMSO. The plate was placed on a shaker for 5 min at 600 RPM. The absorbance was measured at 570 nm using a plate reader (Enspire, Perkin Elmer GmbH, Germany).

### 5.8. Live-cell imaging and direct current stimulation

Seeded devices were placed in an incubated inverted microscope (Zeiss Axio Observer with Definite Focus 2) and maintained at 37°C and 5 % CO_2_. Phase-contrast images were acquired every 3 minutes for the mono-culture and 10 minutes for the co-culture devices using a 10X or 5X objective, respectively.

Constant monophasic DC stimulation (1 to 15 μA for the mono-culture device and 50 μA for the co-culture device) was carried out using a potentiostat/galvanostat (PGSTAT204, Metrohm, Autolab). The current densities used for mono-culture experiments were 0.05 to 0.75 A m^−2^; for the co-culture experiments, it was 0.28 A m^−2^.

#### 5.8.1. Threshold determination

For threshold determination, device 1 (μ1) was employed. After cell seeding, a train of stimulation phases of increasing currents was applied in monophasic direction with 30 minutes pause between the increments. The stimulation time *t*_*s*_ was adjusted for each cell type individually. The parameters used in this study are listed for each cell type in Table 2.

**Table 2:** Parameters for threshold determination and polarity reversal in the single-channel microfluidic device μ1.

|  | threshold determination |  |
| --- | --- | --- |
|  | keratinocytes (K) | fibroblasts (F) |
| $t_s$ [min] | 120 | 180 |
| $EF$ [$\text{mV mm}^{-1}$] | 25 – 50 – 100 – 200 | 50 – 100 – 200 – 400 |
|  | polarity reversal |  |
|  | keratinocytes (K) | fibroblasts (F) |
| $t_s$ [min] | 120 | 180 |
| $EF$ [$\text{mV mm}^{-1}$] | 200 | 400 |

#### 5.8.2. Polarity reversal

For polarity reversal studies, device 1 (μ1) was employed. After cell seeding, the cells were stimulated in alternating directions, starting with a positive current and then a negative one, with 30 minutes pause between the switches. The stimulation time *t*_*s*_ was adjusted for each cell type individually. The parameters used in this study are listed for each cell type in Table 2.

#### 5.8.3. Cellular separation

For cellular separation studies, device 2 (μ2) was utilized. Cells were stimulated for *t*_*s*_ *=* 12 hours at a constant current of *I* = 50 μA. Different electric fields were achieved within the microfluidic device due to its design with branching electrical resistance.

### 5.9. Cell tracking

Time-lapse images were pre-processed with FIJI (ImageJ) to improve contrast, sharpen, and align the frames to the direction of the applied EF. Subsequently, cell tracking was done with the CellTracker toolbox for MATLAB (MathWorks, MA, USA) developed by Piccinini et al.^61^ For each EF tested, as well as for the control experiments, 50 cells were chosen. For the co-culture experiments, 25 keratinocytes and 25 fibroblasts were tracked. The cells were chosen randomly across biological replicates and tracked manually, leaving out any cell that underwent mitosis or left the field of view. The tracked cells were analyzed and plotted in MATLAB to determine their electrotactic parameters:

- **Cell movement:** cell position in x and y were determined at each time point by the distance and angle from the origin and were plotted with their origin being the x-y position at *t* = 0.
- **Directedness (cosθ):** the angle **θ** was defined as the angle between the EF (x-axis) and the migration vector 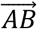 with *A* being the cell position at *t* = 0 and *B* the position at each subsequent time point. We defined the position of the cathode **(-)** to the right and the anode **(+)** to the left of the image. A directedness of 0 equals movement perpendicular to the EF, whereas a value of **+1** represents cathodic and **-1** anodic migration, respectively.
- **Average velocity:** determined by the distance traveled by each cell between frames, divided by the time elapsed

### 5.10. Statistical Analysis

For each experiment, three biological replicates were done under the same conditions. The tracked cells for each replicate were chosen randomly from different locations within the microfluidic devices, and then the data from each position and replicate were pooled to have at least 16 ± 1 cells per replicate to reach 50 total. A Student’s t-test was used with a 99 % confidence interval was used to assess the significance of the differences between the directedness and velocity of the non-stimulated and stimulated cells.

## Data availability

The datasets generated during and/or analysed during the current study are available from the corresponding author on reasonable request.

## Author Contributions

**JL, SS**, and **MA** conceived the project and designed the experiments. **JL** designed and fabricated the mono-culture microfluidic devices and electrodes. **SS** designed and fabricated the co-culture microfluidic devices and electrodes. **NJ** and **AS** performed cell culture and seeding of devices. **SS** performed all FEA simulations, the co-culture media experiments and metabolic assays. **JL** and **SS** performed the mono-culture and co-culture DC stimulation experiments, respectively. **JL** performed all cell tracking, data analysis and visualization. **JL** and **SS** equally co-wrote the manuscript. **JL** prepared figures 1-3 & 5. **SS** prepared figures 4 & 6. All authors reviewed the manuscript. **MA** provided scientific advice/guidance, funding, and supervision for all.

## Acknowledgments

**MA, JL, SS, NJ, & AS:** European Research Council (ERC; 759655; SPEEDER) under the European Union’s Horizon 2020 Research and Innovation program. **MA**: Freiburg Institute for Advanced Studies (FRIAS) and Brainlinks-BrainTools which are funded by the German Research Foundation (DFG; EXC 1086) and is currently funded by the Federal Ministry of Economics, Science and Arts of Baden Württemberg within the sustainability program for projects of the excellence initiative. We would like to thank Mrs. Ute Riede for her immeasurable help with supporting cell culture maintenance and seeding protocols, as well as Mr. Lukas Matter and Ms. Malgorzata Skorupa for their fruitful discussions about the co-culture media and MTT assay experiments.

